# Genomic characterization of the adolescent idiopathic scoliosis associated transcriptome and regulome

**DOI:** 10.1101/2020.03.02.973735

**Authors:** Nadja Makki, Jingjing Zhao, Zhaoyang Liu, Walter L. Eckalbar, Aki Ushiki, Anas M. Khanshour, Zhuoxi Wu, Jonathan Rios, Ryan S. Gray, Carol A. Wise, Nadav Ahituv

## Abstract

Adolescent idiopathic scoliosis (AIS), a sideways curvature of the spine, is the most common pediatric musculoskeletal disorder, affecting ∼3% of the population worldwide. However, its genetic bases and tissues of origin remain largely unknown. Several genome-wide association studies (GWAS) have implicated nucleotide variants in noncoding sequences that control genes with important roles in cartilage, muscle, bone, connective tissue and intervertebral discs (IVDs) as drivers of AIS susceptibility. Here, we set out to define the expression of AIS-associated genes and active regulatory elements by performing RNA-seq and ChIP-seq against H3K27ac in these tissues in mouse and human. Our study highlights genetic pathways involving AIS-associated loci that regulate chondrogenesis, IVD development and connective tissue maintenance and homeostasis. In addition, we identify thousands of putative AIS-associated regulatory elements which may orchestrate tissue-specific expression in musculoskeletal tissues of the spine. Quantification of enhancer activity of several candidate regulatory elements from our study identifies three functional enhancers carrying AIS-associated GWAS SNPs at the *ADGRG6* and *BNC2* loci. Our findings provide a novel genome-wide catalog of AIS-relevant genes and regulatory elements and aid in the identification of novel targets for AIS causality and treatment.

## Introduction

Adolescent idiopathic scoliosis (AIS) is a lateral spinal curvature with no obvious etiology, that manifests during the adolescent growth spurt (1,2). This condition affects around 3% of school-aged children worldwide. The only treatments to prevent curve progression are restrictive bracing or corrective surgery, the latter often requiring extensive postoperative pain management. Hospital costs of operative treatment alone exceed 1 billion USD annually (3). Currently, the genetic basis, the underlying biological mechanisms and the tissues of origin of AIS remain largely unknown. Thus, there are no reliable methods to predict susceptibility to AIS or medications to prevent curve progression. Therefore, it is crucial to gain insight into disease mechanisms and pathogenesis in order to devise new methods to diagnose and treat AIS.

Genome-wide association studies (GWAS) identified at least thirteen genetic loci underlying AIS susceptibility (4–12), all of which reside within noncoding regions of the genome. These results suggest that nucleotide changes in gene regulatory sequences such as enhancers could be potential drivers of AIS susceptibility (8,9). These regulatory elements are thought to control genes with roles in chondrocytes: *ADGRG6* (also called *GPR126*) (5,13), *SOX6, CDH13* (4), *BNC2* (8), *SOX9* (7); intervertebral discs (IVDs): *ADGRG6, PAX1* (9), *SOX9*; muscle: *LBX1* (6,11), *PAX1, BNC2, SOX6, PAX3* (12); bone: *BNC2*; and connective tissue: *FBN1, FBN2* (14), *COL11A2* (15). It is worth noting, that AIS-associated variants are not necessarily causal, but rather point to disease candidates that are in linkage disequilibrium. For the majority of tissues implicated in AIS susceptibility, we currently do not have publicly-available (e.g. ENCODE) genome-wide maps of candidate regulatory elements and also lack transcriptomic data of the genes they could be regulating. Although multiple risk factors are now understood to affect AIS susceptibility, the interplay of these factors in genetic networks and potential common pathways that could be affected have not been defined.

To obtain a genomic understanding of AIS-associated genetic networks and the regulatory elements that control them, we performed RNA-seq and chromatin immunoprecipitation followed by sequencing (ChIP-seq) for the active chromatin mark histone H3 lysine 27 acetylation (H3K27ac) (16,17) on six relevant tissues/cell types: human cartilage, bone and muscle and mouse IVDs, chondrocytes and connective tissue. Our systematic genome-wide analysis establishes an extensive catalog of genes, genetic pathways and active regulatory elements in AIS-relevant tissues. To demonstrate the power of these datasets, we identified gene expression profiles and active regulatory elements for all known AIS GWAS loci in the six examined tissues. We identified putative regulatory elements overlapping AIS-associated SNPs, as well as genetic networks around AIS-associated genes. By performing enhancer assays in chondrocytes, we functionally validated three novel regulatory elements harboring AIS-associated SNPs, two at the *ADGRG6* locus and one at the *BNC2* locus. Our transcriptional and epigenetic profiling of AIS relevant tissues provides a basis for additional functional follow up studies to investigate the pathogenic mechanisms underlying AIS susceptibility.

## Results

### Expression profiling of AIS relevant tissues identifies AIS-associated gene regulatory networks

To systematically identify genes expressed in AIS-relevant tissues, we carried out RNA-seq on human cartilage, muscle and bone and on mouse IVDs, connective tissue and chondrocytes (as we were not able to obtain equivalent human tissues) (Fig 1). The above tissues were chosen as they are implicated in AIS-susceptibility through GWAS and animal studies (4–13). RNA was collected from three biological replicates of each tissue, reverse transcribed and sequenced. These analyses revealed a list of genes expressed in each of the six tissues (we identified over 4,300 genes per tissue), as well as genes that are expressed in multiple tissues (Fig 2A and Table 1). Analysis of AIS associated genes via GWAS found many of these genes to be expressed in at least one of the examined tissues (Table 1). For example, *Adgrg6* expression was detected in mouse chondrocytes, *Kcnj2* in mouse connective tissue and *EPHA4* in human bone. We also identified AIS-associated transcription factors that are known to be critical regulators of chondrogenesis in each of the tissues, such as *SOX9* and *PAX1* in cartilage and IVDs.

**Table 1.**
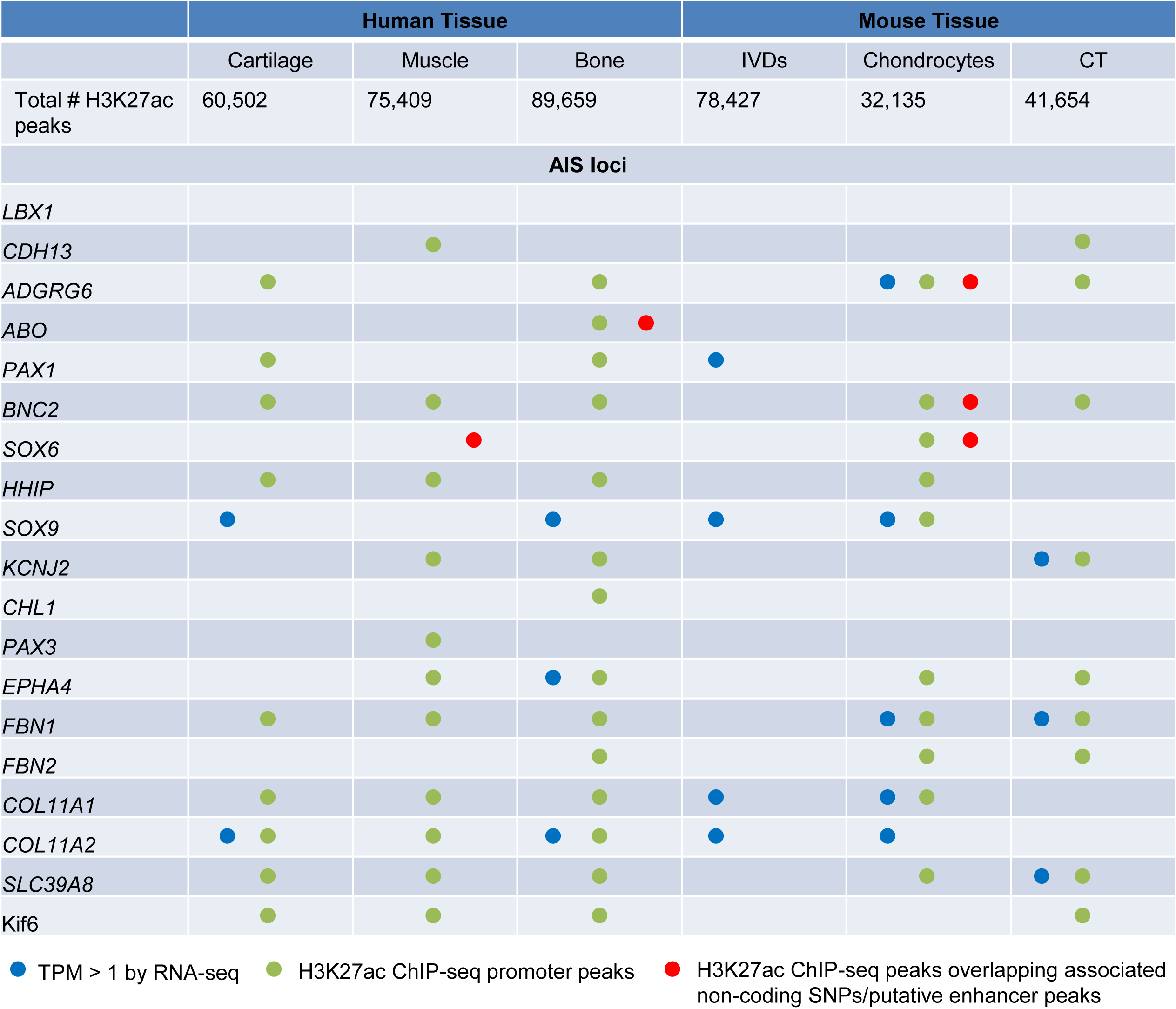
Overview of RNA-seq and ChIP-seq results for AIS GWAS loci. TPM = transcripts per kilobase million.

**Figure 1.**
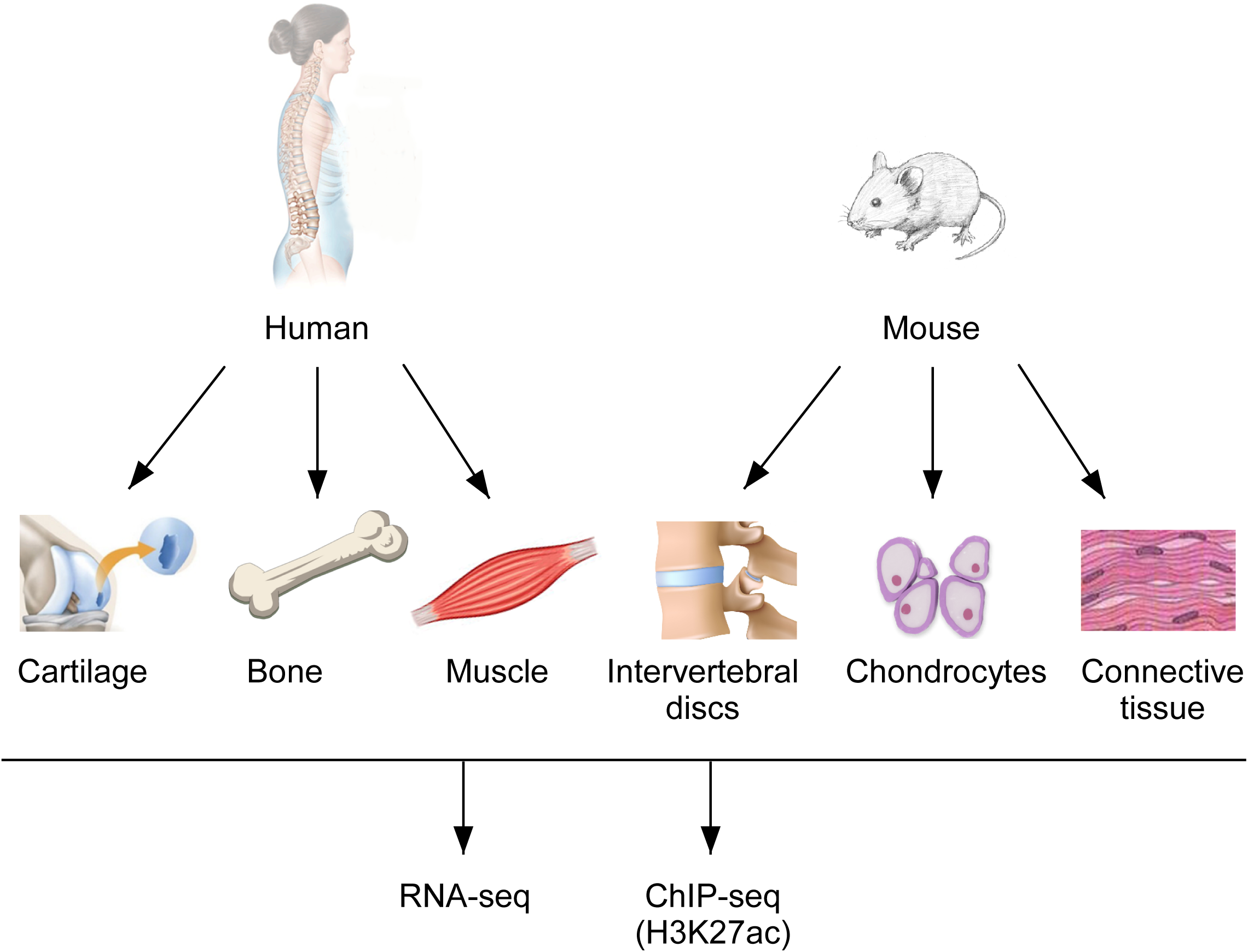
Schematic outline of the study. Cartilage, bone and muscle were isolated from three human individuals and intervertebral discs, chondrocytes and connective tissue were isolated from mice. All tissues were subjected to RNA-seq and H3K27ac ChIP-seq to identify gene expression profiles and tissue-specific regulatory elements.

**Figure 2.**
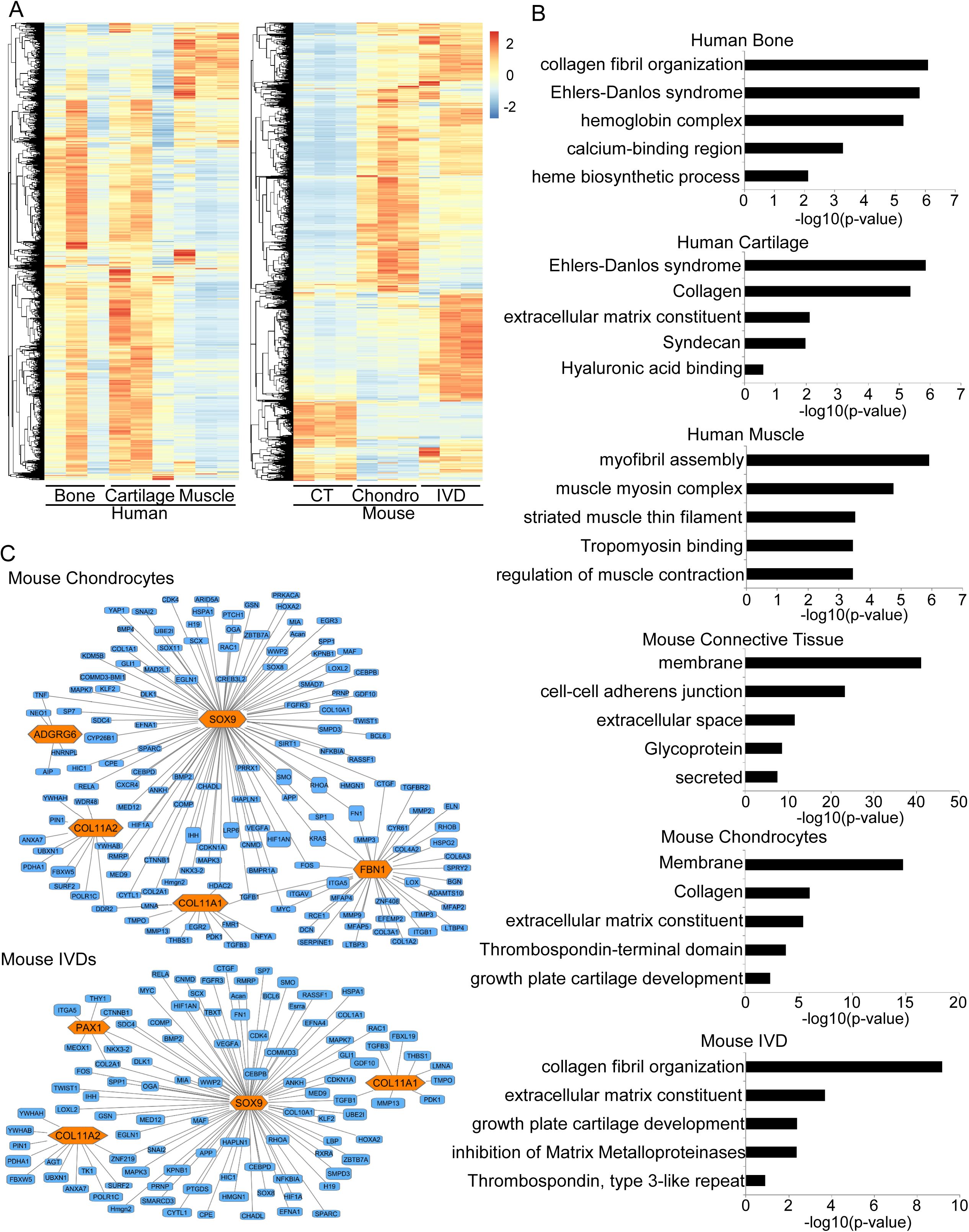
RNA-seq gene expression profiling of AIS relevant tissues and pathway analysis of AIS associated genes. **(A)** Heat maps of gene expression profiles of the six examined tissues. Connective tissue (CT), chondrocytes (Chondro), intervertebral discs (IVD). **(B)** Top gene ontology terms as determined by DAVID (18). **(C)** Gene regulatory networks for AIS-associated loci (shown in orange) as identified using Ingenuity Pathway Analysis.

We next set out to characterize enriched pathways and genetic networks around AIS-associated genes in each tissue. Functional annotation clustering using the Database for Annotation, Visualization and Integrated Discovery (DAVID) (18) found an enrichment of gene ontology (GO) categories for each tissue. For example, chondrocytes are enriched for GO categories related to collagen (p-value = 9.8E-7), extracellular matrix structural constituent (p-value = 4.3E-6) and growth plate cartilage development (p-value = 4.6E-3) (Fig 2B). Further analysis of our tissue-specific datasets using Ingenuity Pathway Analysis (IPA) (19) revealed novel genetic networks up- and down-stream of AIS-associated genes as well as interactions between multiple AIS genes in specific tissues (Fig 2C). For example, tumor necrosis factor-alpha (TNF-alpha) was shown to be a potent inhibitor of *SOX9* in cartilage (20) and *ADGRG6* expression was reported to be decreased when *SOX9* is deleted (21). These analyses highlight genetic networks and putative molecular pathways involved in AIS pathogenesis.

### ChIP-seq identifies AIS-associated regulatory elements

To identify active regulatory elements in AIS relevant tissues, we performed ChIP-seq for H3K27ac on all six tissues (Fig 1). Each tissue was processed using two biological replicates, chromatin was cross linked, immunoprecipitated and DNA sequenced. We found over 30,000 peaks in each tissue (Table 1). We also identified numerous peaks that are shared between two or more tissues. For example, 5,152 peaks were shared between mouse chondrocytes and IVDs (Fig 3A). Using the Genomic Regions Enrichment of Annotations Tool (GREAT; (22)), we observed tissue-specific enrichment of peak regions nearby genes belonging to several GO categories. For example, in chondrocytes we observed a significant association with genes involved in chondrodystrophy (p-value = 1.8E-18), decreased skin tensile strength (p-value = 1.5E-16) and absent somites (p-value = 1.3E-14). In connective tissue, we saw an enrichment with genes involved in joint laxity (p-value = 8.9E-26), vertebral compression fractures (p-value = 1.3183E-8) and premature skin wrinkling (p-value = 1.2589E-7). These results highlight putative mechanisms that might contribute to the pathogenesis of AIS.

**Figure 3.**
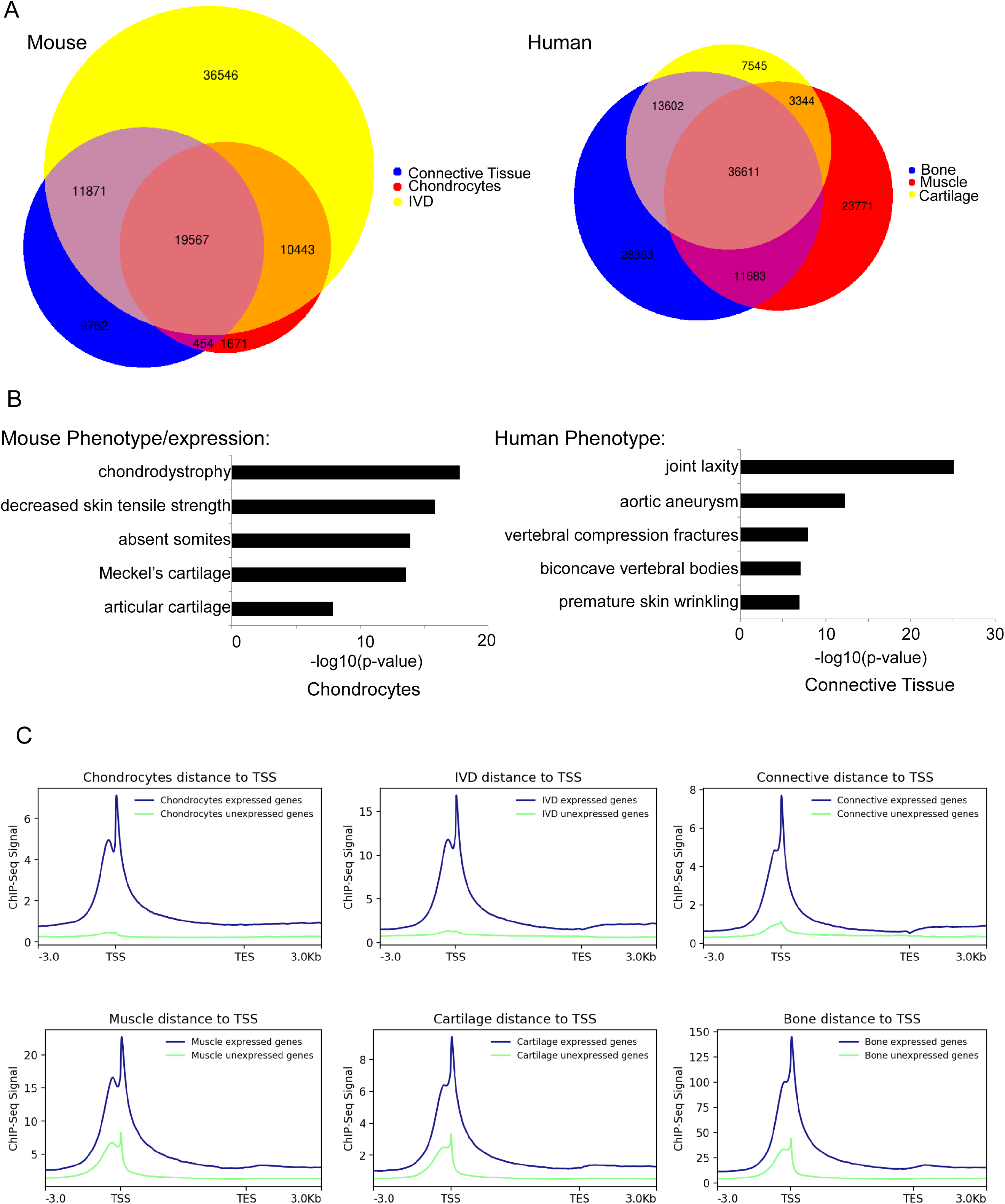
ChIP-seq identifies active regulatory elements in AIS-associated tissues. **(A)** Venn diagram showing the overlap between H3K27ac peaks in mouse and human tissues. **(B)** Top mouse phenotype and gene expression, according to the Mouse Genome Informatics (MGI), and human phenotype term enrichment for mouse chondrocytes and human connective tissue respectively according to GREAT (22). **(C)** H3K27ac peaks show higher correlation around the transcription start site (TSS) of genes that are expressed in the RNA-seq from the same tissue.

In order to test for the correlation between H3K27ac peaks and actively expressed genes in each tissue, as determined by ChIP-seq and RNA-seq respectively, we grouped genes in each tissue into either expressed (transcripts per kilobase million (TPM) ≥ 1) or not expressed and determined for each group the correlation to ChIP-seq signal and the distance to the transcriptional start site (TSS). We found a positive correlation between the actively expressed genes and regions having H3K27ac signals for each tissue (Fig 3C). These findings corroborate the overlapping gene ontology term enrichments found for both our RNA-seq and ChIP-seq genes and peaks (Fig 2B and Fig 3B). GO terms related to cartilage development and extracellular matrix constituent were found for both active genes and H3K27ac peaks. We also examined the correlation between our RNA-seq and ChIP-seq data specifically for the AIS-associated loci and found that many associated genes that are expressed (as determined by RNA-seq) have H3K27ac peaks around the TSS (Table 1).

### Identification and functional characterization of novel AIS-associated enhancers

One major challenge in the functional follow up of GWAS is to assign function to noncoding variants, in particular as the lead SNP is not necessarily pathogenic. We thus set out to determine whether AIS-associated SNPs identified in previous GWAS overlap with putative regulatory elements identified in our ChIP-seq experiments. For these analyses, we identified all SNPs in linkage disequilibrium (r2>=0.8) with lead AIS GWAS SNPs and overlaid them with H3K27ac peaks for each of the six tissues. We identified tissue-specific H3K27ac peaks that overlap GWAS SNPs for several of the AIS-associated loci (Table 1). These sequences represent putative regulatory elements that could be affected by the disease associated SNPs.

To functionally validate some of these sequences, we focused our analysis on H3K27ac peaks identified in mouse chondrocytes, as chondrocytes were implicated as a major cell type underlying AIS susceptibility (13,21,23) and existing cell lines can be easily transfected for *in vitro* validation experiments. We identified chondrocyte-specific H3K27ac peaks overlapping AIS-associated SNPs at the *ADGRG6* (Fig 4A) and *BNC2* loci. We selected seven sequences at the *ADGRG6* locus and three at the *BNC2* locus to test for enhancer activity using luciferase reporter assays. Sequences were cloned into an enhancer assay vector (pGL4.23; Promega), which contains a minimal promoter followed by a luciferase reporter gene. As a positive control, we used pGL4.13 (Promega) with an SV40 early enhancer (pGL4.13; Promega) and the pGL4.23 empty vector as a negative control. Out of the ten assayed sequences, three showed significant luciferase activity in a human chondrocyte cell line (SW1353) (Fig 4B and S1 Fig). Two of these functional enhancers are located in the *ADGRG6* locus with one sequence (*ADGRG6_4*) residing in the fourth intron and the second (*ADGRG6_6*) in the 3’UTR of this gene (Fig 4B). The third sequence is located in the third intron of *BNC2* overlapping an alternative promoter (*BNC2_2*) (S1 Fig). In sum, we discovered three novel AIS-associated regulatory elements that could affect gene expression levels and contribute to disease susceptibility.

**Figure 4.**
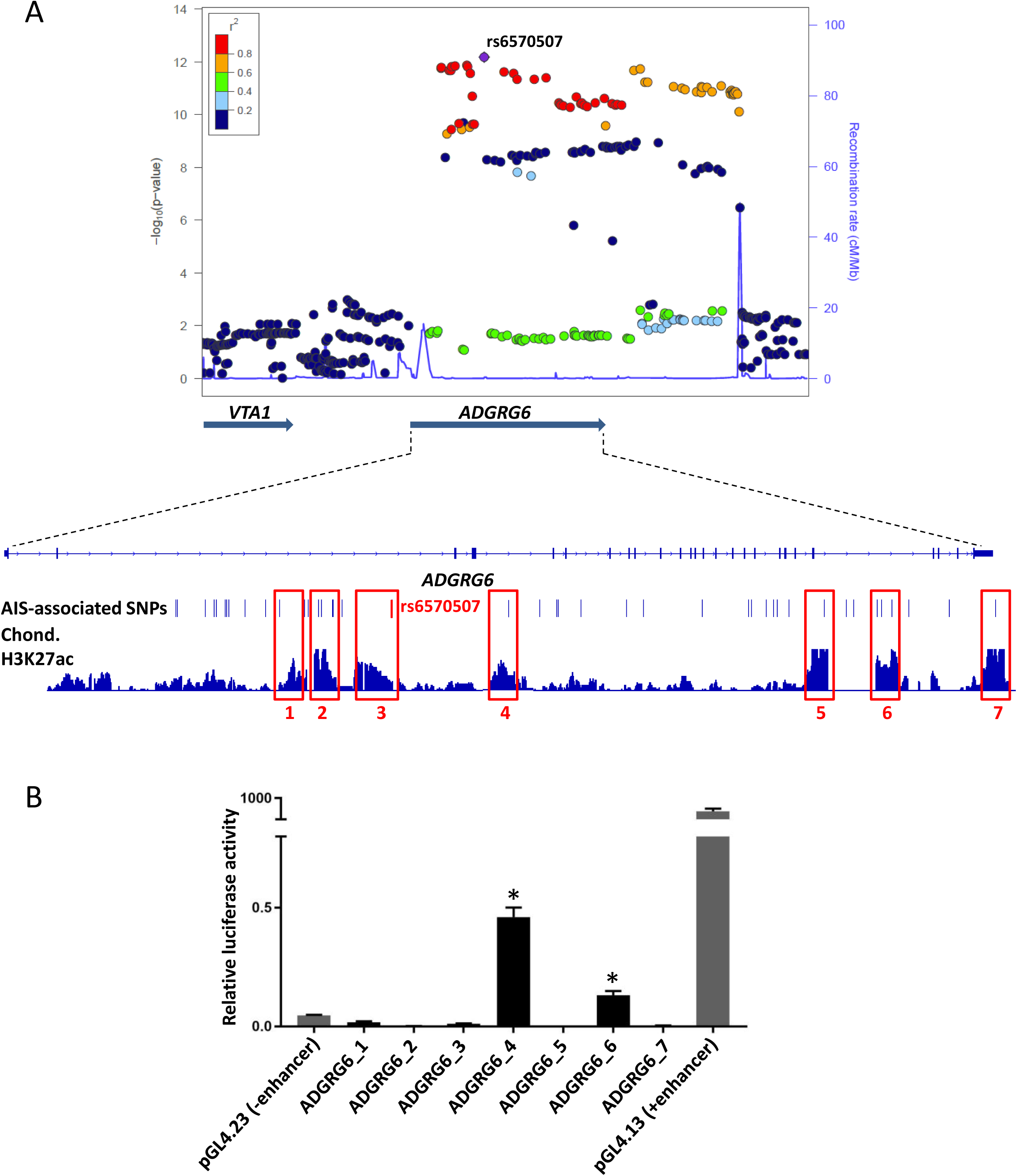
Identification and characterization of AIS-associated enhancers at the *ADGRG6* locus. **(A)** Integrative genomic viewer snapshot and Locuszoom plot for the *ADGRG6* locus highlighting the overlap of AIS-associated SNPs with H3K27ac ChIP-seq peaks. The Locuszoom plot is indexed by the lead SNP rs6570507(4). The colors of the spots reflect the degree of linkage disequilibrium (LD) with the index SNP measured by r^2^. **(B)** Luciferase assays in human SW1353 cells identify two novel enhancers at the *ADGRG6* locus (*= P < 0.02).

## Discussion

In this study, we set out to identify genes, genetic networks and regulatory elements underlying AIS susceptibility in a genome-wide manner. We performed RNA-seq and ChIP-seq on six AIS relevant tissues: IVDs, chondrocytes, connective tissue, cartilage, bone and muscle. Our RNA-seq results allowed us to define enriched pathways and biological processes in these tissues. We also identified transcriptional profiles and novel genetic networks of important AIS genes. For example, the genetic network surrounding *ADGRG6*, an important AIS susceptibility gene (4,5,13), showed interactions with several other genes that have been implicated in AIS, with one of these genes being *SOX9*, a master regulator of chondrogenesis (24). This interaction is supported by direct observation that loss of *Adgrg6* in the IVD is associated with reduced SOX9 expression in cartilaginous endplate and annulus fibrosus regions of the IVD in mouse (21). Thus our analyses allowed us to identify genetic networks and predict interactions between several AIS genes, which will assist future studies on the intricate interplay of these loci in driving susceptibly to this complex disease.

Our H3K27ac ChIP-seq datasets revealed thousands of novel putative enhancers in the six tissues and provided exceptional candidate sequences for subsequent studies to probe the functional relevance of AIS-associated non-coding SNPs. Using GREAT, we observed tissue-specific enrichment and a significant association with genes involved in human and mouse phenotypes such as chondrodystrophy and vertebral compression fractures that might contribute to the pathogenesis of scoliosis. We also found a strong overlap between active genes as shown by RNA-seq and active ChIP-seq chromatin marks by both genome-wide comparisons as well as by examining specific AIS loci, demonstrating the correlation between our datasets.

One major focus of post GWAS studies is to assign function to disease-associated non-coding SNPs. As a first step in this direction, we generated a database of tissue-specific regulatory regions in AIS-relevant tissues. We used these data to identify novel putative enhancers overlapping with associated SNPs at several AIS-associated GWAS loci: *LBX1, CDH13, GPR126, ABO, PAX1, BNC2, SOX6, HHIP, SOX9*/*KCNJ2, CHL1, PAX3, AJAP1*/*EPHA4* and *BCL2* (4–12). These putative AIS-associated enhancers are a valuable resource for future functional and mechanistic follow up studies. As a great majority of risk factors underlying AIS remain undiscovered, future GWAS, whole-genome sequencing and family studies will uncover many additional AIS loci, which can then also be analyzed using our genomic datasets.

As chondrocytes were shown to be an important cell type underlying AIS susceptibility (13,21,23), we functionally tested putative chondrocyte enhancers that overlap AIS-associated SNPs using enhancer assays. We identified three novel functional enhancer sequences that encompass SNPs in LD with AIS GWAS SNPs. One enhancer is located in the fourth intron and one in the 3’UTR of *ADGRG6* and a third enhancer is located in the third intron of *BNC2* overlapping an alternative promoter. The *ADGRG6* and *BNC2* loci were found to be associated with AIS both via GWAS and animal studies (4,5,8,13). Additional work on how these loci could affect their expression and subsequent function would be of interest. *ADGRG6* in particular represents an exciting locus as it encodes for a G-protein coupled receptor and is therefore a potential ‘druggable’ target. Further investigation is needed to dissect the regulation and interactions of *ADGRG6* and other AIS-associated loci as well as shared mechanisms that lead to AIS susceptibility. Our study provides the first comprehensive genome-wide survey of transcriptional and epigenetic profiles of AIS-relevant tissues and highlights novel AIS-associated regulatory elements and genetic networks, enabling the identification of new candidates for the diagnosis and treatment of AIS.

## Methods

### Human and mouse tissue samples

All human tissue samples were collected at the Texas Scottish Rite Hospital for Children in Dallas and approved by an Institutional Review Board. Facets and muscle were collected from the thoracolumbar spine of AIS patients ages 11-17 during spinal surgery and cartilage was dissected off the bone. Tissues were snap-frozen for RNA-seq or fixed in 1% formaldehyde when used for ChIP-seq (see below). We isolated mouse connective tissue from 8–10 week old CD1 mice (Charles River), intervertebral discs (IVDs) from the thoracic and lumbar spine of postnatal day (P) 20 mice and costal chondrocytes from the anterior rib cage and sternum of P 2– 4 mice. Chondrocytes were isolated as previously described (13). Briefly, rib cages were dissected, soft tissues were removed and rib cages digested at 37°C for 1 hour in 2 mg/ml pronase and then washed in phosphate-buffered saline (PBS). Rib cages were then further digested for 1 hour using 3 mg/ml collagenase D in Dulbecco’s Modified Eagle Medium at 37°C and 5% CO_2_ followed by 3 PBS washes to remove digested tissue. Remaining cartilage was then further digested for 4–6 hours in 3 mg/ml collagenase D and then filtered through a 45 μm cell strainer to obtain chondrocytes.

### RNA isolation and sequencing

Each tissue was processed using three biological replicates. Each human replicate consisted of a tissue from one patient. For mouse chondrocytes, we pooled tissue from 6-10 mice per replicates, for connective tissue we pooled 4 mice and for IVDs we isolated 20-24 IVDs from 2 mice as one replicate. RNA was isolated from chondrocytes using the RNeasy Mini Kit and from connective tissue using the RNeasy Fibrous Tissue Mini Kit (Qiagen). All other tissues were first homogenized using Lysing Matrix S, 50 х 2mL Tubes (Fisher, 116925050) in TRIzol Reagent (Invitrogen, 15596026) on the Tissue Lyser II. Total RNA was then cleaned up with the Direct-zol RNA miniprep kit (Zymo Research, Z2070). RNA quantity and purity were analyzed on a Bioanalyzer 2100 and RNA 6000 Nano LabChip Kit (Agilent). Total RNA was subjected to isolate Poly (A) mRNA with poly-T oligo attached magnetic beads (Invitrogen). RNA fragments were reverse-transcribed to create the final cDNA libraries following the NEBNext® Ultra™ RNA Library Prep Kit (Illumina) and paired-end sequencing 150bp was performed on a Novaseq 6000. Sequence quality was verified using FastQC (25). Raw sequencing reads were mapped to the mouse (GRCm38/m10) or human (GRCh37/hg19) genome (hg19) using STAR (26).

### Chromatin immunoprecipitation-sequencing (ChIP-seq)

Each tissue was processed using two biological replicates and samples were pooled in the same manner described above for RNA sequencing. Tissues were cross linked using 1% formaldehyde by standard techniques (27). ChIP was performed using an antibody against H3K27ac (Millipore, Burlington, MA 05-1334) using the LowCell# ChIP kit (Diagenode). Illumina sequencing libraries were generated using the Accel-NGS 2S Plus DNA Library Kit (Swift Biosciences). Single-end 50bp sequencing was done on a HiSeq 4000 and computational analyses were performed using BWA (28) and MACS2 (29).

### Luciferase reporter assays

Enhancer candidate sequences (S2 Table) were selected based on having an H3K27ac peak in mouse chondrocytes and overlapping SNPs in linkage disequilibrium (LD) with an AIS-associated GWAS lead SNP. The SNPs in LD were determined using ldlink (30) on data from the 1000 Genomes Project (31,32) in the HapMap Caucasian (CEU) population. Enhancer candidate sequences were PCR amplified from human genomic DNA, cloned into a pGL4.23 enhancer assay vector (Promega), and sequence-verified. Empty pGL4.23 was used as negative control and pGL4.13 (Promega) with an SV40 early enhancer as a positive control. Luciferase assays for enhancer candidates were carried out in chondrocyte cell lines (human SW1353 cells (ATCC® HTB-94™). SW1353 cells were grown in DMEM (Invitrogen). SW1353 media was supplemented with 1% penicillin/streptomycin and 10% and 5% FBS respectively. Forty-eight hours before transfection, 60,000 cells were plated out in 96-well plates and were grown up to 90% confluency. Cells were transfected with 130 ng of the assayed plasmid and 16 ng of pGL4.73 [hRluc/SV40] (Promega) containing Renilla luciferase to correct for transfection efficiency, using X-tremeGENE (Roche) according to the manufacturer’s protocol. Transfections were performed in triplicates. Forty-eight hours after transfection, cells were lysed, and luciferase activity was measured using the Dual-Luciferase Reporter Assay Kit (Promega). Measurements were performed on a GloMax 96-microplate luminometer (Promega).

## Figure Legends

**Supplemental Figure 1.**
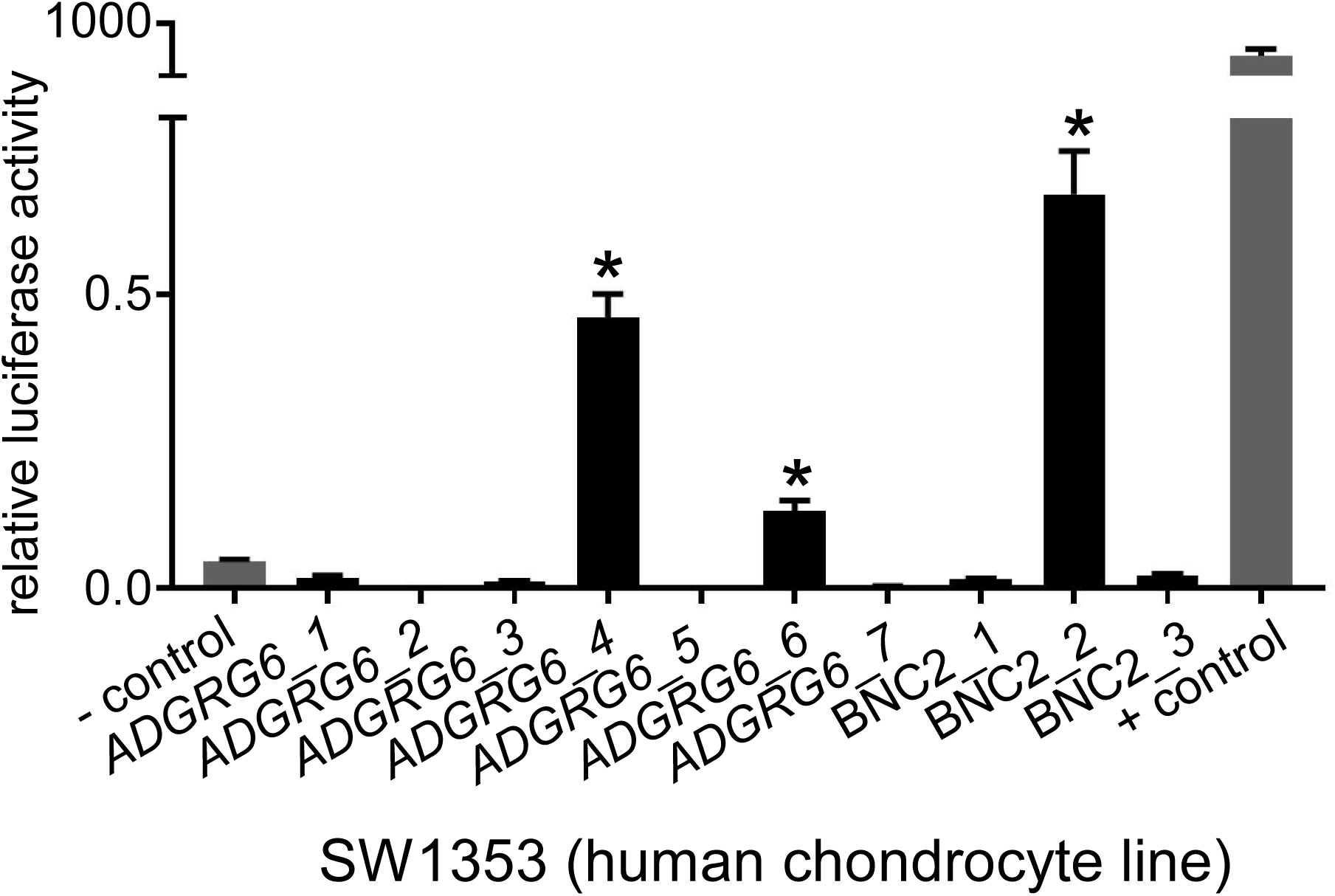
Enhancer assays for the *ADGRG6* and *BNC2* loci in human SW1353 cells.

**Supplemental Table 1.**
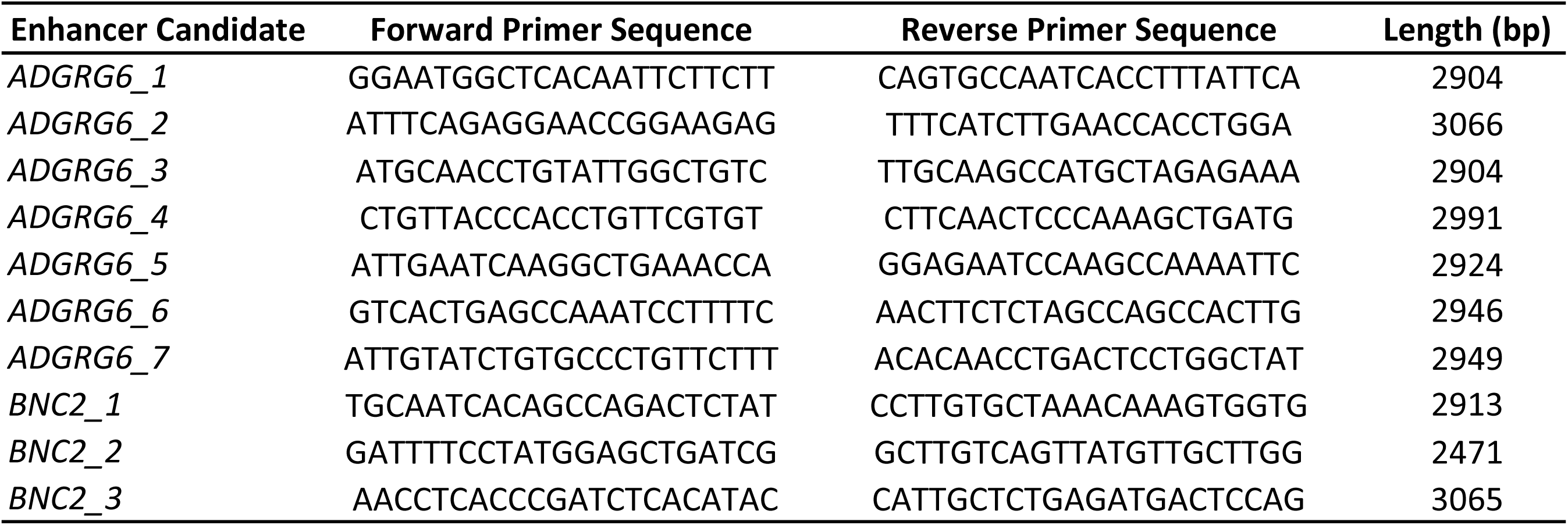
Primer sequences for enhancer candidate regions at the *ADGRG6*and *BNC2* loci.

## Author contributions

NM and NA conceived the experiments. NM performed the experiments. JZ, NM, WLE and ZW performed RNA-seq and ChIP-seq analysis. AU helped with ChIP-seq samples. AMK compiled AIS SNPs. ZL, RSG, JR and CAW provided tissues, expertise, and feedback. NM and NA wrote the manuscript.

## Acknowledgements

We would like to thank Fumitaka Inoue for his help with sequence sample submission and Anna Williams and Nandina Paria for their help with human tissue collection. Research in this publication was supported in part by the National Institutes of Child and Human Development grant number 1P01HD084387 (N.A. and C.A.W.) and the Arthritis and Musculoskeletal and Skin Diseases R01AR072009-01 (R.S.G.) and F32AR073648 (Z.L.).

